# Combined 2-deoxy-D-glucose and metformin selectively depletes glioblastoma cells earlier than primary human astrocytes

**DOI:** 10.64898/2026.11.23.768727

**Authors:** B Jewett, C Groberg, L Zaddack, A Carter, O Harris-Norman, B Page, P Bogues, R Kellar

## Abstract

Glioblastoma (GBM) cells depend heavily on glycolysis for growth, which makes combined inhibition of glycolysis and mitochondrial respiration an attractive treatment strategy. Whether normal glia tolerate such a combination, and whether the effect persists at the low oxygen levels of brain tissue, is less clear. We treated U-118 MG GBM cells and primary human cortical astrocytes with 2-deoxy-D-glucose (2-DG, 4 mM), metformin (2 mM), or both, under normoxia and 2% O2, and measured cell number (CyQUANT) and metabolic activity (PrestoBlue) in the same wells. At day 3, in separate experiments for each cell type, the combination reduced U-118 MG cell number to 55.0 ± 8.0% of vehicle under normoxia and 73.7 ± 14.0% under 2% O2, while astrocyte cell number was not detectably reduced (103.8 ± 7.6% and 96.6 ± 20.3%; mean ± SD of three plates). Across both oxygen levels the difference between cell types was 35.8 points (95% CI 16.5 to 55.1; p = 0.002). Metformin alone had little effect on cell number (88% to 121% of vehicle), but it potentiated 2-DG: across all plates, adding metformin lowered cell number a further 1.77-fold in U-118 MG (95% CI 1.44 to 2.18) and 1.16-fold in astrocytes (95% CI 1.07 to 1.26). At day 5, with both cell types treated and read together, U-118 MG cell number had fallen to 20.7 ± 4.6% (normoxia) and 20.8 ± 6.8% (2% O2), and astrocytes retained 51.1 ± 1.5% and 50.4 ± 1.6%, a difference of about 30 points at each oxygen level with no detected interaction between cell type and oxygen level. In a single EdU incorporation experiment, combination-treated U-118 MG cells exhibited minimal EdU incorporation. U-118 MG cells therefore responded earlier and more deeply than astrocytes to 2-DG plus metformin, at both oxygen levels. Astrocytes followed the same course later and to a lesser depth.

## Introduction

Glioblastoma (GBM) is the most common malignant primary brain tumor [1]. With radiotherapy plus temozolomide, median survival in the trial that set the standard of care was 14.6 months [2]. GBM cells proliferate rapidly, and rapid proliferation carries a high demand for energy and biosynthetic precursors. Like many proliferating tumor cells, GBM cells rely on glycolysis even when oxygen is available [3]. GBM also contains hypoxic regions, which contribute to resistance to radiotherapy because oxygen makes radiation-induced DNA damage permanent [4].

2-Deoxy-D-glucose (2-DG) is a glucose analog that undergoes phosphorylation by hexokinase, resulting in the accumulation of 2-DG-6-phosphate and subsequent inhibition of glycolysis [5]. Metformin is a first-line drug for type 2 diabetes. It inhibits mitochondrial complex I, part of the electron transport chain, and has been studied as an anticancer agent [6]. Combining the two should limit both major routes of ATP production. The combination of 2-DG and metformin depleted ATP and reduced viability in cancer cells [7, 8]. It was active across a broad range of cancer cell lines and slowed xenograft growth [8]. In glioma models, the combined treatment reduced proliferation of pediatric glioma lines [9] and suppressed viability, stemness, and invasion of GBM tumorspheres, with a potential survival benefit in an orthotopic model [10].

Two questions matter for any metabolic combination aimed at brain tumors. The first is whether normal glia tolerate the treatment; the second is whether the effect holds at the oxygen levels cells encounter in the brain. Standard cell culture conditions typically maintain oxygen near atmospheric levels, substantially higher than the physiological oxygen levels found in brain tissue (roughly 2% to 9%), which decrease further within the tumor [11]. GBM cells that invade past the resection margin and seed recurrence sit in this low-oxygen tissue, so a result obtained only at atmospheric oxygen may not transfer. The glioma studies of the combination did not include normal glial cells or low-oxygen conditions [9, 10]. Evidence exists for each drug alone. Primary rat astrocytes were resistant to the antiproliferative and proapoptotic action of metformin on glioma cells [12]. Metformin (10 mM) reduced viable cell number in several glioma lines at 5% O2, with a similar tendency reported at 1% O2 [11]. Hypoxia was reported to protect primary GBM cells from 2-DG [13]. Here we compare U-118 MG GBM cells and primary human cortical astrocytes treated with 2-DG, metformin, or both, under normoxia and 2% O2, using cell number, estimated from DNA content, as the primary readout.

## Methods

### Cell culture

U-118 MG human GBM cells (ATCC HTB-15, lot 70045102) were used at passages 2 to 9 and maintained on uncoated tissue-culture plastic in high-glucose DMEM (Gibco 10566016) with 10% fetal bovine serum (Gibco A3160401) at 37 °C and 5% CO2. The ATCC certificate of analysis for this lot reports a human STR profile meeting ATCC specification and no mycoplasma detected by Hoechst stain, agar culture, or PCR. The cells used here were not re-authenticated in the laboratory; STR validation is pending. Primary human cortical astrocytes (ScienCell, catalog 1800) were used at passage 2 and maintained in Astrocyte Medium (Gibco A1261301; DMEM with 1% N-2 supplement and 10% fetal bovine serum) and cultured on tissue-culture plastic coated with Matrigel basement membrane matrix (Corning 354234, diluted 1:6) for both day 3 and day 5 experiments. Both media are based on high-glucose DMEM (4.5 g/L glucose).

### Drug treatment and oxygen conditions

2-DG (MilliporeSigma D8375) and metformin hydrochloride (MilliporeSigma D150959) were prepared in culture medium. Both cell types were seeded in 96-well plates at 10,000 cells per well for plates read at day 1 or day 3 and 7,500 cells per well for plates read at day 5 (lowered so that wells did not reach confluence before the read), and treated with vehicle, 4 mM 2-DG, 2 mM metformin, or 4 mM 2-DG plus 2 mM metformin. Each experimental condition was represented by six wells per plate. Each plate was seeded from a separate flask of cells, with the three flasks within each experimental group originating from three independent cell thaws, so the three plates served as independent biological replicates (n = 3), while the six wells within each plate served as technical replicates. All plates treated on the same day were dosed from the same drug preparation. Normoxia and 2% O2 plates of the same replicate were seeded and treated together.

Plates read at day 1 or day 3 received drug once without a medium change. For plates read at day 5, medium with drug was refreshed at day 3. Plates were kept at 37 °C and 5% CO2 either in a standard humidified incubator at ambient oxygen (normoxia) or in a hypoxia incubator at 2% O2. Hypoxic incubator oxygen was monitored with a pumped gas analyzer (FD-600, Forensics Detectors) and remained at 2 ± 0.5%. The laboratory is in Flagstaff, Arizona, at approximately 2,100 m elevation, so ambient oxygen partial pressure is lower than at sea level.

### PrestoBlue and CyQUANT assays

PrestoBlue (Invitrogen A13261) and CyQUANT Direct (Invitrogen C35011) assays were read sequentially on the same wells on a Synergy HT microplate reader (BioTek) after incubation with PrestoBlue reagent for 20 min and CyQUANT reagent for 1 h (PrestoBlue 530/25 nm excitation, 590/35 nm emission; CyQUANT 485/20 nm excitation, 528/20 nm emission). Each plate carried two blank wells. Signal was blank-subtracted and expressed as a percentage of the mean of the vehicle wells on the same plate. CyQUANT Direct reports the DNA content of live cells and was used as the estimate of cell number.

PrestoBlue reports the total reducing activity of the well and was used as the measure of metabolic activity. Their ratio (blank-corrected PrestoBlue divided by blank-corrected CyQUANT, the metabolic index) is reported as metabolic activity per cell. One vehicle well with a CyQUANT signal below one third of its condition median was excluded (day 5, 2% O2, plate 1, well F8).

Plates were read at day 3 and at day 5 in separate experiments with the same doses, plate layout, and three plates per oxygen condition for each cell type. At day 3, the astrocyte and U-118 MG plates were run as separate experiments about one month apart. At day 5, the plates of both cell types were treated on the same day and read together.

### Dose matrices

For range-finding of drug doses, cells were treated for 24 h in a checkerboard of 2-DG (0, 1, 2, 3, 4, 5 mM) and metformin (0, 1, 2, 2.5, 3, 4 mM), with each dose pair in duplicate wells per plate and read as above. Each plate also carried three to six additional zero-dose wells outside the matrix in row A or in rows A and H. Signal was expressed as a percentage of the mean of all zero-dose wells on the plate, the outer-row wells together with the two inside the matrix (for the U-118 MG normoxia plate, PrestoBlue was referenced to the two wells inside the matrix). On the astrocyte plates analyzed here, the outer-row zero-dose wells read lower than those inside the matrix (PrestoBlue by 18 to 22%, CyQUANT by 8 to 21%). Because 24 h is early for a change in cell number, the metabolic index was used to compare U-118 MG and astrocytes across the matrix at 2% O2. Matrices were also run under normoxia for both cell types. The difference between cell types was largest at 4 mM 2-DG plus 2 mM metformin, and this pair was carried forward. Dose selection was exploratory: the pair was chosen because it gave the largest difference between cell types on these plates, which can overstate the difference seen later. One of two astrocyte plates at 2% O2 was excluded from the matrix analysis because of a row artifact: the wells at 4 mM 2-DG read about 40% above the neighboring rows in both assays, and the wells at 5 mM 2-DG differed by about 50% between the left and right halves of the plate. One of two U-118 MG plates under normoxia was excluded because CyQUANT signal in three wells at 5 mM 2-DG was at the level of the blank. Results with the excluded astrocyte plate put back are given in the Results.

Interaction between the two drugs was assessed on the same plates by Bliss independence. On each plate, the metabolic index at every dose pair was expressed as a fraction of the mean of all zero-dose wells. The expected fraction for a combination was the product of the fractions for 2-DG alone and metformin alone at the same doses, and the Bliss excess was the expected minus the observed value in percentage points. A positive excess means a larger reduction than independent action predicts.

### EdU incorporation and cleaved caspase-3 staining

Staining followed a written protocol fixed before the first replicate (version 1.4). U-118 MG cells were seeded at 20,000 cells per well in 8-well chamber slides (ibidi µ-Slide 8 Well, ibiTreat) and treated the next day with vehicle, 4 mM 2-DG, 2 mM metformin, or the combination for 3 days without a medium change, with duplicate wells per condition. EdU (10 µM) was added for the final 2 h before fixation in 4% paraformaldehyde for 15 min. Fixed slides were sealed with Parafilm and stored in a humidified bag at 4 °C for one week before staining. Cells were permeabilized in 0.5% Triton X-100 for 20 min. EdU was detected with the Click-iT Plus EdU Alexa Fluor 647 Imaging Kit (Invitrogen C10640). After blocking in 5% normal goat serum with 0.3% Triton X-100 for 60 min, cells were incubated for 24 h at 4 °C with cleaved caspase-3 (Asp175) rabbit monoclonal antibody (clone 5A1E, Cell Signaling Technology 9664; 1:400), then for 3 h with Alexa Fluor 568 goat anti-rabbit secondary antibody (A11011, lot 2013083, diluted 1:200). Nuclei were stained with Hoechst 33342. A control slide processed in the same session carried an apoptosis positive control (UV exposure for 1 min), a DMSO vehicle well, a proliferation-arrest control (palbociclib, 1 µM), a well without EdU, wells without primary antibody, and an additional combination-treated well.

Wells were imaged on a Leica TCS SPE confocal microscope (10x/0.30 objective; 405, 561 and 635 nm excitation; one 4 x 4 tile scan per well, single plane).

Nuclei were segmented from the Hoechst channel with fixed parameters. Following the protocol, a nucleus was scored EdU-positive when its mean EdU-channel signal exceeded the mean plus 3 SD of nuclei in the no-EdU well. One well that received no EdU (a metformin duplicate) was excluded.

### Statistics

The plate was the unit of replication. Each plate was an independent biological replicate seeded from a separate flask and a separate cell thaw (n = 3 per group unless stated, each the mean of six wells). Within a cell type, the combination was compared with vehicle by a one-sample t-test against 100% and with each single agent by a paired t-test across plates. U-118 MG and astrocytes were compared by Welch’s t-test. Because normoxia and 2% O2 plates of each replicate were run together, comparisons between oxygen levels within a cell type used paired t-tests across replicates. P values were adjusted by the Holm method within each family of six comparisons. CyQUANT was the primary readout. The contribution of metformin to the combination was expressed per plate as a potentiation ratio, the cell number with 2-DG alone divided by the cell number with the combination (a ratio of 1 means metformin added nothing). Ratios were analyzed on the log scale, tested against 1 with a one-sample t-test, compared between cell types with Welch’s t-test, and are reported as geometric means with 95% confidence intervals, pooled across the day 3 and day 5 experiments and for each day. At day 3, the effect of cell type on combination-treated cell number was also estimated across both oxygen levels with a linear model (cell type and oxygen as factors, plates as units), and at day 5 by two-way ANOVA with the same factors. Bliss excess from the day 1 matrices is reported per plate without a test, because each group has one or two plates. Analyses used Python 3.15 with SciPy 1.18. Values are mean ± SD.

## Results

### Dose selection from day 1 matrices

U-118 MG cells and astrocytes were treated for 24 h with a 6 x 6 matrix of 2-DG and metformin at 2% O2 (Figure 1). Metabolic activity per cell fell with increasing 2-DG in both cell types, more in U-118 MG. Of the 36 dose pairs, the difference between cell types was largest at 4 mM 2-DG plus 2 mM metformin, where the metabolic index was 66% of control in U-118 MG and 87% in astrocytes (a difference of 20 points). Neighboring pairs gave similar differences (19 to 20 points at 3 or 4 mM 2-DG with 2.5 mM metformin). The size of the difference depended on the reference wells: with only the two zero-dose wells inside the matrix as the reference it was 11 points at 4 mM plus 2 mM, still the largest of the 36. With the excluded astrocyte plate put back, the difference at this pair was 13 points and ranked third of 36. The 4 mM plus 2 mM pair was used for all later experiments. This comparison rested on two U-118 MG plates and one astrocyte plate, with two wells per dose pair, and is descriptive. PrestoBlue and CyQUANT values for each matrix are given in Figure S1.

**Figure 1.**
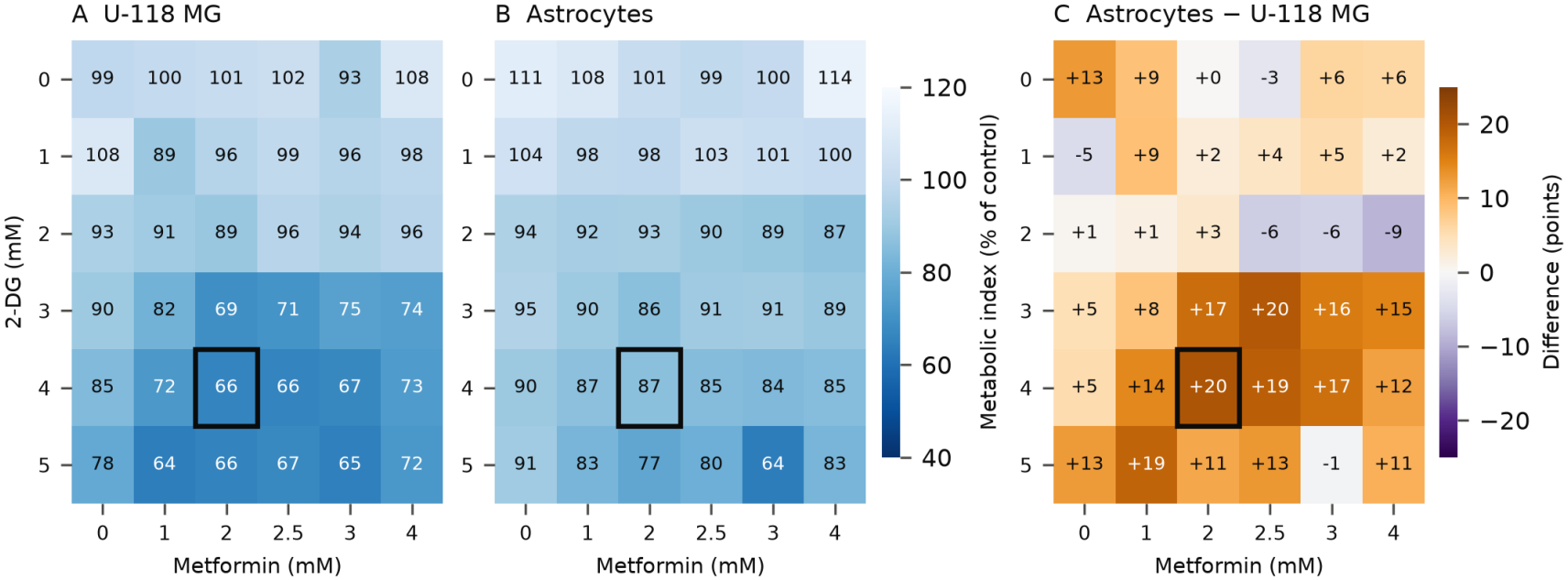
Dose selection from day 1 matrices. Metabolic index (PrestoBlue divided by CyQUANT) after 24 h of treatment with 2-DG and metformin at 2% O2 in (A) U-118 MG cells and (B) astrocytes, as a percentage of the mean of all zero-dose wells on the plate (the two inside the matrix and the additional wells in the outer rows). (C) Difference between cell types (astrocytes minus U-118 MG, percentage points). U-118 MG values are the mean of two plates and astrocyte values come from one plate (a second astrocyte plate was excluded for a row artifact; see Methods), with two wells per dose pair per plate. The outlined cell marks 4 mM 2-DG plus 2 mM metformin, used in all later experiments.

### At day 1 the Bliss excess at 4 mM plus 2 mM was larger in U-118 MG than in astrocytes at 2% O2

The same matrices were used to ask whether the two drugs acted independently (Figure S2). At 4 mM 2-DG plus 2 mM metformin and 2% O2, the metabolic index in U-118 MG was 23.5 and 14.8 points below the Bliss prediction on the two plates, against 4.0 points in astrocytes (one plate), consistent with synergy in U-118 MG at this dose pair. Across the whole matrix, the two cell types were closer. The excess was positive at 24 and 17 of the 25 dose pairs in U-118 MG (17 on both plates; mean excess 15.0 and 9.0 points) and at all 25 pairs in astrocytes (mean 10.5 points). Under normoxia, the excess in astrocytes was positive at 25 and 24 of the 25 pairs on two plates (mean 6.6 points on each; 3.1 and 2.8 points at 4 mM plus 2 mM). In U-118 MG under normoxia, on the one usable plate, the excess at 4 mM plus 2 mM was -8.8 points. It was positive at 10 of the 25 pairs, five of them in the 3 mM metformin column, where values of 36 to 114 points suggest a plate artifact. For PrestoBlue signal alone, the excess in U-118 MG at 2% O2 was -9.9 and 4.6 points at 4 mM plus 2 mM. On the excluded astrocyte plate, at 2% O2 the excess at 4mM plus 2 mM was 12.9 points. Each single-agent value rested on two wells per plate, and the excess depends on the reference wells: on the astrocyte plates, the outer-row zero-dose wells read lower than those inside the matrix (Methods), which raises every fraction and the expected value with it. These matrices are exploratory.

### At day 3 the combination reduced U-118 MG cell number and did not detectably reduce astrocyte cell number

U-118 MG cells and astrocytes were treated in separate experiments and read at day 3 (Figure 2A, B). In astrocytes, cell number with the combination was 103.8 ± 7.6% of vehicle under normoxia and 96.6 ± 20.3% under 2% O2 (n = 3 plates each). Neither differed from vehicle (p = 0.48 and 0.80). 2-DG alone and metformin alone also left astrocyte cell number at or above 90% of vehicle (112.8 ± 0.7% and 93.8 ± 0.5% under normoxia; 122.5 ± 6.3% and 97.5 ± 2.2% under 2% O2).

**Figure 2.**
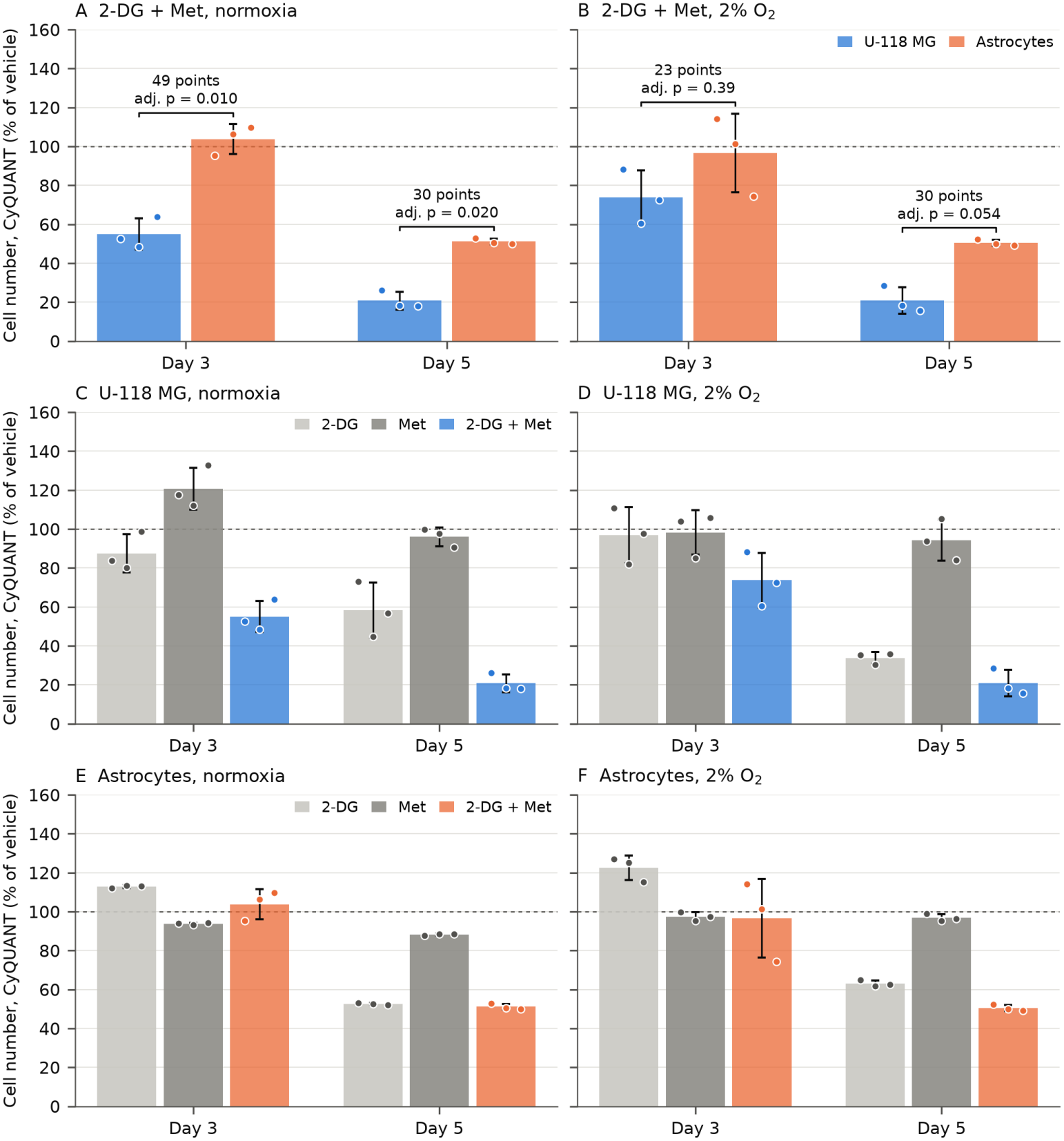
The combination reduced U-118 MG cell number at day 3 and day 5, and astrocyte cell number only by day 5. Cell number by CyQUANT, as percent of the vehicle wells on the same plate. (A, B) Cells treated with 4 mM 2-DG plus 2 mM metformin (2-DG + Met), U-118 MG next to astrocytes, under normoxia and 2% O2. Brackets give the difference between cell types (astrocytes minus U-118 MG, percentage points) and its Holm-adjusted p value. (C, D) U-118 MG and (E, F) astrocytes treated with 2-DG, metformin (Met), or both, under normoxia and 2% O2. Within each panel the day 3 and day 5 groups are separate experiments (10,000 and 7,500 cells seeded per well; drug refreshed at day 3 in the day 5 experiment), so they are not joined as a time course. At day 3 the U-118 MG and astrocyte plates were run in separate experiments about one month apart; at day 5 they were treated on the same day. Bars show the mean of plates, points the individual plates (independent biological replicates seeded from separate flasks, six wells each), error bars SD. n = 3 plates per group. One vehicle well was excluded (U-118 MG, day 5, 2% O2, plate 1, well F8; see Methods). The dashed line marks vehicle (100%).

In U-118 MG, the combination reduced cell number to 55.0 ± 8.0% of vehicle under normoxia. This was lower than 2-DG alone (87.5 ± 9.8%, adjusted p = 0.005) and metformin alone (120.7 ± 10.7%, adjusted p = 0.003). Under 2% O2, the combination gave 73.7 ± 14.0%, lower than 2-DG alone (96.8 ± 14.4%, adjusted p = 0.009); the differences from metformin alone (98.3 ± 11.4%) and from vehicle did not reach significance after adjustment (both adjusted p = 0.083).

Across both oxygen levels, combination-treated astrocytes retained 35.8 points more cell number than U-118 MG (95% CI 16.5 to 55.1; p = 0.002; 6 plates per cell type), with no significant dependence on oxygen (interaction p = 0.13). Taken separately, the difference was 48.8 points under normoxia (95% CI 31.0 to 66.6; p = 0.002, adjusted p = 0.010) and 22.9 points under 2% O2 (95% CI -18.7 to 64.4; p = 0.19, adjusted p = 0.39; Figure 3A). The interval under 2% O2 was wide, so these data do not rule out an effect of oxygen on the difference between cell types.

**Figure 3.**
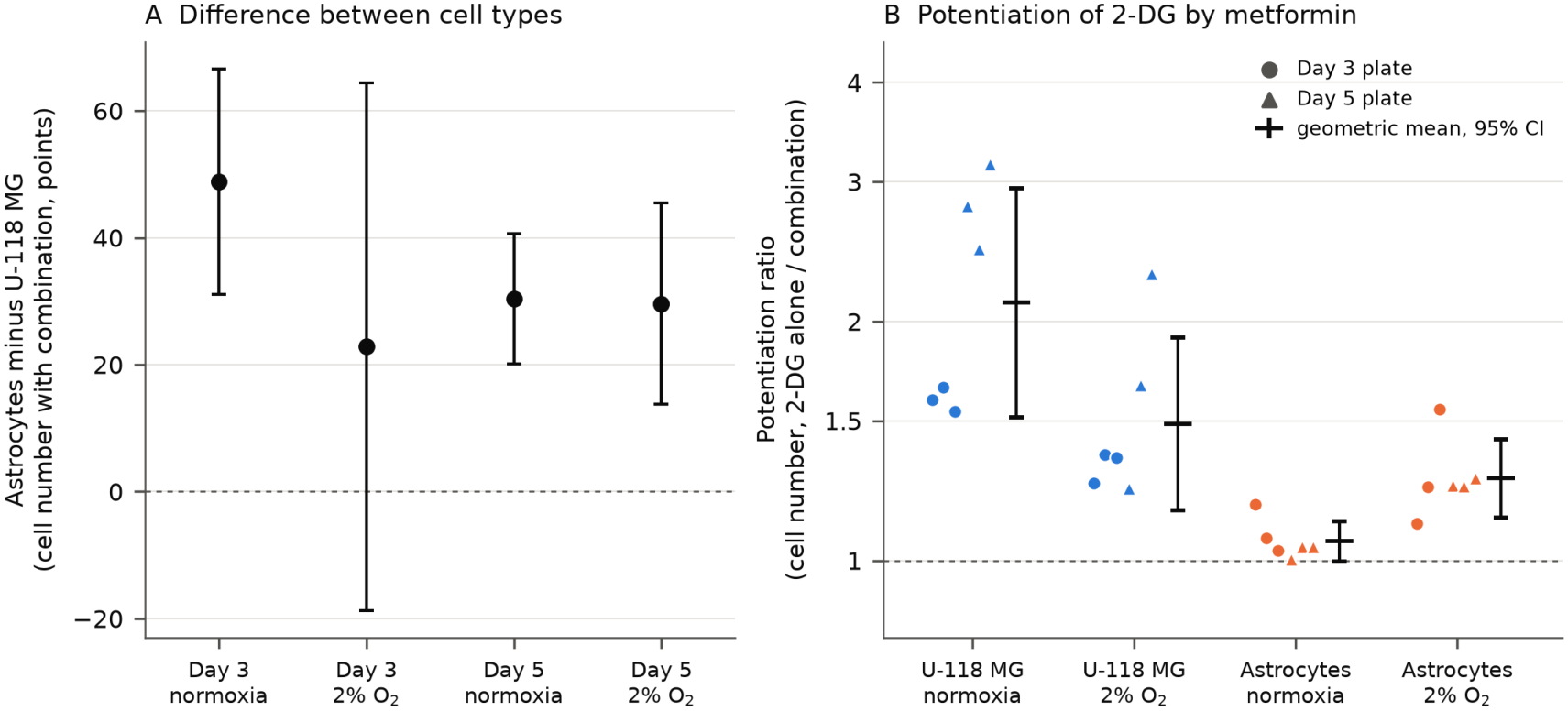
Difference between cell types and potentiation of 2-DG by metformin. (A) Cell number with the combination in astrocytes minus U-118 MG, in percentage points of vehicle, at day 3 and day 5 under normoxia and 2% O2. Points are the difference of group means (three plates per cell type) and error bars the 95% confidence interval. The dashed line marks no difference. (B) Potentiation ratio for each plate: cell number with 2-DG alone divided by cell number with the combination, on a log scale. Circles are day 3 plates and triangles day 5 plates (six plates per group). Horizontal marks and error bars are the geometric mean and 95% confidence interval. The dashed line marks a ratio of 1, where metformin adds nothing.

Metabolic activity was lower with the combination in both cell types. PrestoBlue signal fell to 37.9 ± 11.1% (normoxia) and 54.5 ± 2.4% (2% O2) of vehicle in U-118 MG (adjusted p = 0.031 and 0.005). In astrocytes, it was 73.4 ± 19.9% and 60.4 ± 19.1% (Figure S3); these values were lower than with metformin alone (adjusted p = 0.005 and 0.048) but did not differ significantly from vehicle (p = 0.15 and 0.07). At day 3, the combination therefore did not detectably reduce astrocyte cell number, with a tendency toward lower metabolic activity.

### By day 5 the combination reduced cell number in both cell types, with greater loss in U-118 MG

At day 5, U-118 MG cells and astrocytes were treated on the same day and read together (Figure 2). The effect in U-118 MG was larger than at day 3. The combination reduced cell number to 20.7 ± 4.6% under normoxia and 20.8 ± 6.8% under 2% O2 (adjusted p = 0.006 and 0.010 against vehicle). 2-DG alone gave 58.2 ± 14.2% and 33.7 ± 3.0%. The combination was lower than 2-DG alone under normoxia (adjusted p = 0.048); under 2% O2 the same comparison did not reach significance (adjusted p = 0.081). Metformin alone left cell number near vehicle (95.9 ± 4.7% and 94.3 ± 10.6%).

With combination treatment, astrocytes demonstrated a biological response. The combination reduced astrocyte cell number to 51.1 ± 1.5% under normoxia and 50.4 ± 1.6% under 2% O2. 2-DG alone gave 52.5 ± 0.5% and 63.0 ± 1.6%, and metformin alone 88.2 ± 0.5% and 96.8 ± 1.9%. In astrocytes, the combination did not differ from 2-DG alone under normoxia (p = 0.15) and was 12.6 points lower under 2% O2 (adjusted p = 0.003).

Astrocytes still retained more cell number than U-118 MG. The difference with the combination was 30.4 points under normoxia (95% CI 20.1 to 40.6; adjusted p = 0.020) and 29.6 points under 2% O2 (95% CI 13.8 to 45.5; adjusted p = 0.054; Figure 3A). With the excluded vehicle well put back, U-118 MG cell number under 2% O2 was 22.5 ± 9.9% and the difference was 27.8 points (95% CI 4.2 to 51.5). Two-way ANOVA showed an effect of cell type (F(1,8) = 147.8, p < 0.0001) and no interaction with oxygen (F(1,8) = 0.02, p = 0.88); the differences at the two oxygen levels differed by 0.8 points (95% CI -10.6 to 12.2). 2-DG alone separated the cell types only under 2% O2 (33.7% against 63.0%; adjusted p = 0.004).

PrestoBlue followed the same direction (Figure S3): 7.2 ± 2.1% and 15.3 ± 4.2% of vehicle in U-118 MG against 32.4 ± 3.5% and 33.1 ± 3.2% in astrocytes (adjusted p = 0.007 and 0.026). Within U-118 MG, where normoxia and 2% O2 plates were paired by replicate, cell number with the combination did not differ between oxygen levels (paired difference 0.0 points; 95% CI -6.0 to 6.1).

### Metformin potentiated 2-DG, more in U-118 MG than in astrocytes

Metformin alone had little effect on cell number (88% to 121% of vehicle across cell types, times and oxygen levels; the lowest value was 88.2 ± 0.5%, in astrocytes under normoxia at day 5). Its contribution to the combination was measured on each plate as the ratio of cell number with 2-DG alone to cell number with the combination (Figure 3B). In U-118 MG, adding metformin lowered cell number a further 1.77-fold across all 12 plates (95% CI 1.44 to 2.18; p < 0.001): 2.11-fold under normoxia (95% CI 1.51 to 2.95; p = 0.002; 6 plates) and 1.49-fold under 2% O2 (95% CI 1.16 to 1.91; p = 0.010; 6 plates). In astrocytes, the ratio was 1.16 across 12 plates (95% CI 1.07 to 1.26; p = 0.002): 1.06 under normoxia (95% CI 1.00 to 1.12; p = 0.057) and 1.27 under 2% O2 (95% CI 1.13 to 1.42; p = 0.003).

The potentiation was larger in U-118 MG than in astrocytes by a factor of 1.53 overall (95% CI 1.23 to 1.90; p = 0.001). The difference between cell types was clear under normoxia (factor 2.00; 95% CI 1.43 to 2.78; p = 0.003) and was not significant under 2% O2 (factor 1.17; 95% CI 0.91 to 1.51; p = 0.18). Oxygen shifted the potentiation in opposite directions in the two cell types. Paired by replicate, potentiation in U-118 MG was 1.42-fold greater under normoxia than under 2% O2 (95% CI 1.10 to 1.84; p = 0.018; six replicate pairs), and in astrocytes, it was greater under 2% O2 (normoxia to 2% O2 ratio 0.83; 95% CI 0.71 to 0.97; p = 0.030). The ratios above pool the day 3 and day 5 experiments, which were run separately at different seeding densities. By day, the ratio in U-118 MG was 1.45 at day 3 (95% CI 1.29 to 1.63; p < 0.001; 6 plates) and 2.16 at day 5 (95% CI 1.50 to 3.13; p = 0.003; 6 plates). In astrocytes, it was 1.18 at day 3 (95% CI 1.01 to 1.38; p = 0.038) and 1.13 at day 5 (95% CI 1.01 to 1.27; p = 0.037).

### Metabolic activity per cell

The metabolic index was used to separate a change in metabolic activity per cell from a change in the number of cells (Figure S4). Metformin alone left the index between 96% and 108% of vehicle, except at day 3 under normoxia (84.5 ± 18.6% in U-118 MG and 133.1 ± 22.1% in astrocytes). 2-DG alone lowered the index under normoxia (52.2% in U-118 MG and 78.4% in astrocytes at day 5) and left it near or above control in U-118 MG under 2% O2 (106.3%). With the combination, the index in surviving U-118 MG cells at day 5 was 34.3 ± 2.2% under normoxia and 76.4 ± 19.5% under 2% O2, although cell number was the same at the two oxygen levels (about 21%). In astrocytes at day 5, the index with the combination was 63% at both oxygen levels. The combination thus removed a similar fraction of U-118 MG cells at both oxygen levels, while the cells that remained at 2% O2 kept more reducing activity per cell than those that remained under normoxia.

### EdU incorporation and cleaved caspase-3 (replicate 1)

In the two vehicle-treated wells of U-118 MG cells on the treatment slide, 21.0% and 35.3% of nuclei were EdU-positive (188 of 896 and 214 of 606; Figure 4). Three vehicle-treated wells on the control slide, stained in the same session, gave 26.3 to 34.2%. The fraction was 22.4% and 24.2% in the two 2-DG wells and 29.3% in the metformin well, all within the range of the vehicle wells. In the three combination-treated wells, 0 of 153, 7 of 368 and 3 of 132 nuclei were EdU-positive (0%, 1.9% and 2.3%), and nuclear density was 54, 130 and 52 per mm^2^ against 215 to 651 per mm^2^ in vehicle wells. The combination wells resembled the palbociclib arrest control (0.9% EdU-positive). The no-EdU control gave 0.2%.

**Figure 4.**
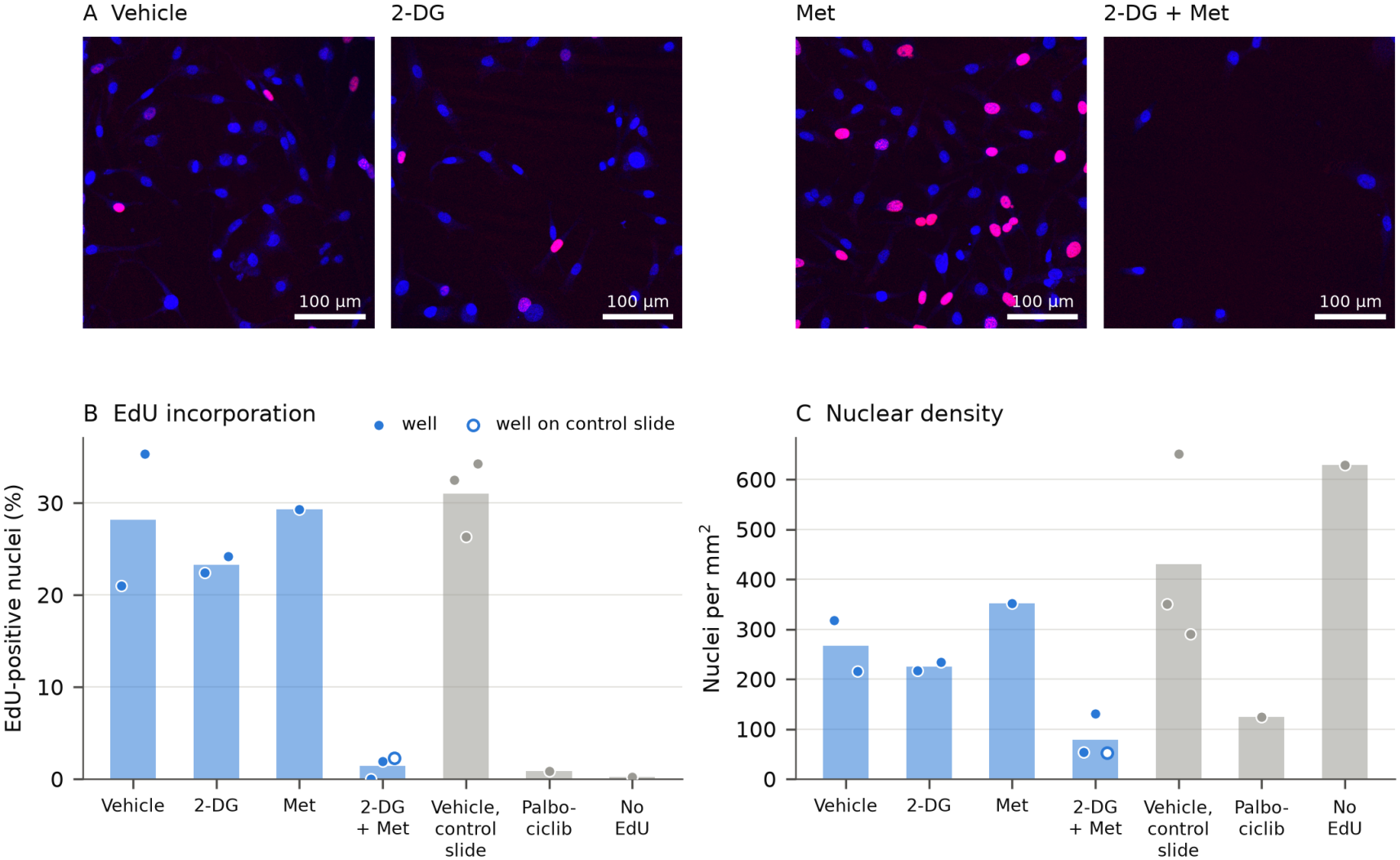
EdU incorporation in U-118 MG cells, replicate 1. (A) Representative fields. Blue, nuclei (Hoechst); magenta, EdU. The cleaved caspase-3 channel is not shown because the stain did not meet its acceptance criterion. Display intensity was doubled equally for all fields. Scale bars, 100 µm. (B) EdU-positive nuclei as a percentage of nuclei counted per well. (C) Nuclei per mm^2^. Each point is one well and bars show the mean of the wells in each group (one well for metformin, palbociclib and the no-EdU control). The open marker is the combination-treated well that sat on the control slide. Grey marks wells from the control slide stained in the same session: three vehicle-treated wells (DMSO vehicle, and two vehicle wells used as antibody controls), palbociclib (1 µM, proliferation arrest), and a vehicle well without EdU.

Cleaved caspase-3 could not be evaluated in this replicate. The UV-treated positive control did not produce a distinct caspase-3-positive population (99th percentile of percell signal 28 of 255, against 29 in the DMSO vehicle well), so the staining did not meet the protocol’s acceptance criterion and no conclusion about apoptosis was drawn. The secondary antibody lot dated from 2021 and is the suspected cause.

Conditions other than vehicle were represented by one to three wells from a single run, so no statistical test was applied.

## Discussion

At day 3, 4 mM 2-DG plus 2 mM metformin decreased U-118 MG cell number by 45% under normoxia and by 26% at 2% O2, and did not detectably reduce astrocyte cell number. By day 5 the combination had reduced U-118 MG cell number by 79% at both oxygen levels and astrocyte cell number by 49 to 50%. U-118 MG cells therefore responded earlier and more deeply than astrocytes. No interaction with oxygen was detected, but with three plates per group, the data cannot rule one out: at day 3, the difference between cell types under 2% O2 had a 95% CI of -18.7 to 64.4 points.

Metformin alone had little effect on cell number, yet it potentiated 2-DG. In U-118 MG, adding metformin lowered cell number an additional 1.8-fold on average, and the potentiation held at both oxygen levels. In astrocytes, the effect was minimal (1.2-fold). Oxygen influenced the effect metformin had in the two cell types. Under normoxia, metformin potentiated 2-DG twice as much in U-118 MG as in astrocytes, where it added almost nothing, so the difference between cell types came largely from metformin. Under 2% O2, the potentiation was more similar in the two cell types (1.5-fold against 1.3-fold) and 2-DG alone separated them by day 5 (33.7% against 63.0%). At day 1, the dose matrices allowed the interaction to be assessed against Bliss independence. At 4 mM plus 2 mM and 2% O2, the excess for metabolic activity per cell was larger in U-118 MG (23.5 and 14.8 points) than in astrocytes (4.0 points), consistent with synergy in U-118 MG at this dose pair. Across the full matrix at 2% O2, the excess was positive in both cell types and of similar size. Under normoxia, U-118 MG demonstrated no synergy at that time point, and the excess in astrocytes was minimal (6.6 points on average on each of two plates). These matrices had one or two plates per group and are exploratory. At day 3 and day 5, a single dose pair was tested, so the effect on cell number is described as potentiation; synergy at those time points would require dose-response data for 2-DG with and without metformin. The metformin concentration used here (2 mM) is below the 8 mM and 5 mM used with 2-DG in earlier glioma studies [9, 10], but it is far above the concentrations reached in patients: most mechanistic studies have used metformin at 10 to 100 times the maximal therapeutic concentration [14]. The present results describe an effect at an in vitro concentration.

Astrocytes were not unaffected. At day 3, the PrestoBlue signal tended to be lower (by 27% and 40%, not significant against vehicle) while cell number was not detectably reduced, and by day 5, cell number had fallen by 49 to 50%. The result was a difference in the timing and depth of the response, and it did not show that the combination is safe for normal glia. It did point to exposure time as a variable worth testing: a shorter exposure may widen the difference between tumor cells and astrocytes.

Reported as a ratio, cell number and metabolic activity from the same wells showed that U-118 MG cells surviving the combination at 2% O2 kept more reducing activity per cell than survivors under normoxia, although the same fraction of cells was lost. The higher reducing activity per cell at 2% O2 may reflect a shift in redox state. PrestoBlue reports cellular reducing capacity and is not specific to mitochondrial respiration.

An earlier report found that hypoxia protected primary GBM cells from the apoptotic effect of 2-DG [13]. Our day 3 results pointed the same way: U-118 MG cell number with 2-DG alone was 96.8% of vehicle at 2% O2 against 87.5% under normoxia, and with the combination 73.7% against 55.0%, although neither difference was significant in paired tests (p = 0.40 and 0.16). By day 5, the direction had reversed for 2-DG alone (33.7% at 2% O2 against 58.2% under normoxia; paired p = 0.07). Cell source and readout differ between the studies (primary cells and apoptosis in that study; an established line and DNA content in ours), and the present data do not fully explain the difference.

Several limitations apply in the current study. Astrocytes were grown on Matrigel-coated vessels in astrocyte medium and U-118 MG cells on uncoated plastic in DMEM with serum, the conditions routinely used for each cell type. Both media are high-glucose DMEM with 10% serum, so glucose supply was matched, but substrate and the N-2 supplement differed between the cell types. Both media contained 25 mM glucose, which competes with 2-DG, so the effect at the lower glucose levels found in tissue may differ.

Only one GBM line and one astrocyte source were tested. The day 3 comparison between cell types was made across two experiments run about one month apart; only the day 5 comparison was made within one experiment. Groups had three plates each, and the day 5 comparison under 2% O2 had an adjusted p of 0.054. The three plates of a group came from separate cell thaws but were dosed from one drug preparation on the same day, so reproducibility across drug preparations is untested. No plate was read at the time of treatment, so growth rates could not be compared; a faster-dividing population loses more percent of vehicle under the same degree of growth inhibition [15], and this may contribute to the difference between cell types. CyQUANT measures the DNA of cells that remain attached and cannot separate cell death, arrest, and detachment. The 2% O2 atmosphere was verified with an oxygen meter, but oxygen at the cell layer and the cellular response to it (for example HIF-1α or CA9) were not measured, and day 5 plates were briefly in room air during drug refresh. The day 1 matrices used for dose selection and for the Bliss analysis had one or two plates per group, with two wells per dose pair, and the dose pair was selected on the outcome that was tested later. The EdU experiment was a single run, and cleaved caspase-3 could not be evaluated from it.

Future experiments will address these limitations by including a second GBM line, a 2-DG dose-response with and without metformin, a hypoxia marker, and replicate EdU and caspase-3 staining.

In summary, 2-DG plus metformin reduced U-118 MG cell number earlier and to a greater extent than astrocyte cell number, under both normoxia and 2% O2, and metformin contributed mainly by potentiating 2-DG. Astrocytes followed the same course later, so the difference between tumor cells and normal glia was one of timing and depth. If that difference holds in additional GBM lines and under matched culture conditions, exposure time becomes a variable that can be set to widen the difference between tumor cells and normal glia. The metformin concentration used here is beyond what systemic dosing reaches, and systemic dosing gives little control over exposure time, so both considerations point toward local, time-limited delivery at the resection margin, where recurrence arises at low oxygen. More broadly, comparing tumor and normal cells side by side, over time and at tissue-level oxygen, is a way to investigate, for any metabolic therapy, whether a therapeutic window exists and when it closes.

## Author contributions

Conceptualization: B.J., P.B. Methodology: B.J., C.G., A.C., O.H.-N., B.P., P.B. Investigation: B.J., C.G., L.Z., A.C., O.H.-N., B.P. Formal analysis: B.J. Visualization: B.J. Data curation: B.J. Funding acquisition: B.J. Project administration: B.J. Resources: R.K. Supervision: B.J., R.K. Writing (original draft): B.J. Writing (review and editing): C.G., A.C., R.K.

## Funding

This work was supported by Hooper Undergraduate Research Awards (2025, 2026) to B.J. from the Office of Undergraduate Research and Creative Activity, Northern Arizona University.

## Competing interests

The authors declare no competing interests.

## Data availability

The data supporting the findings of this study are available from the corresponding author on request. A complete data package will be deposited in a public repository.

## Notes

### Competing Interest Statement

The authors have declared no competing interest.

