## Supplementary Figures S1-S4 for "Combined 2-deoxy-D-glucose and metformin selectively depletes glioblastoma cells earlier than primary human astrocytes"

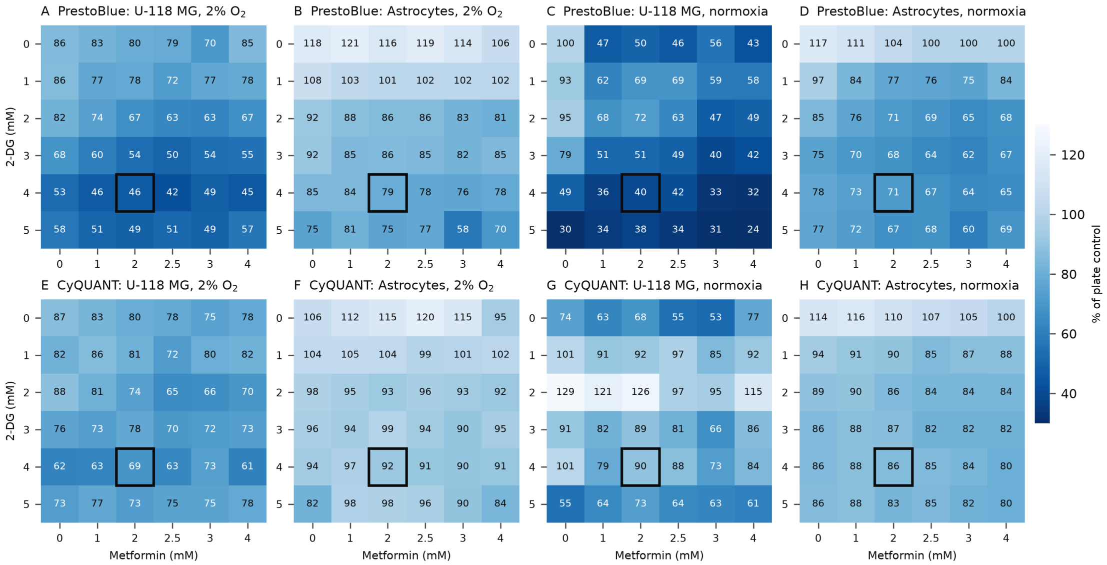


**Figure S1. Day 1 dose matrices, PrestoBlue and CyQUANT.** Signal after 24 h of treatment as a percentage of the mean of all zero-dose wells on the plate, averaged over two plates per group (one plate for astrocytes at 2% O2 and for U-118 MG under normoxia). The outlined cell marks 4 mM 2-DG plus 2 mM metformin.

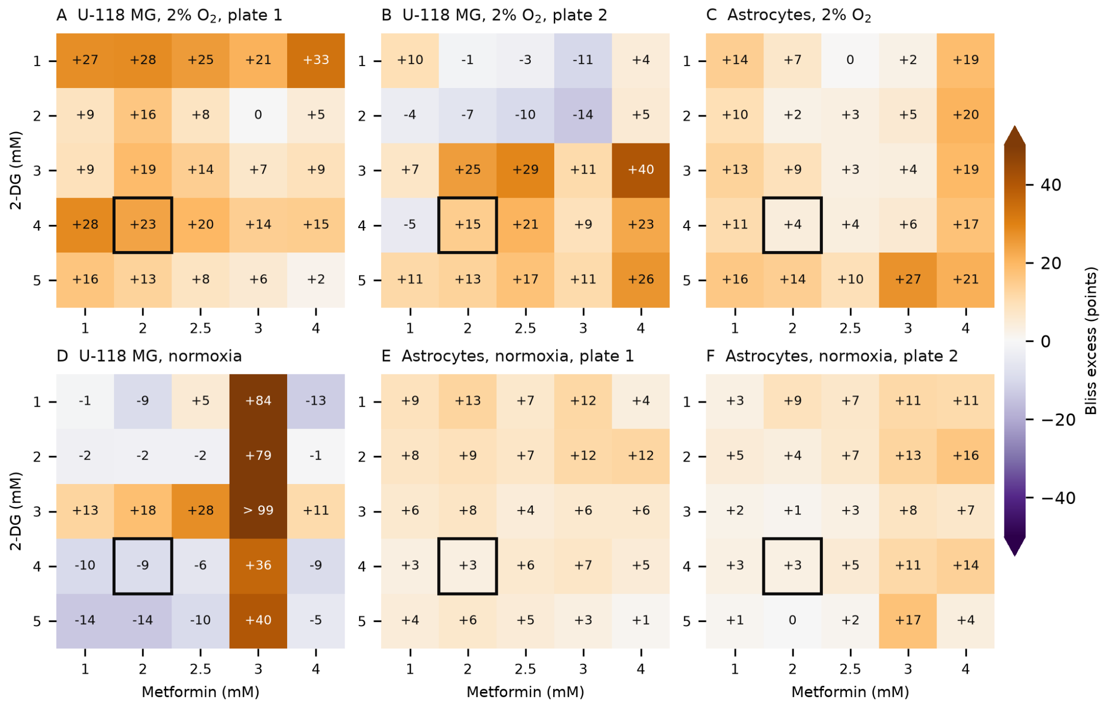


**Figure S2. Bliss excess for the metabolic index at day 1.** Expected minus observed metabolic index, in percentage points, for each combination of 2-DG and metformin after 24 h. Each value was first expressed as a fraction of the mean of all zero-dose wells on the plate, and the expected value was calculated by Bliss independence from the single-agent wells of the same plate. Positive values mean a larger reduction than independent action predicts. (A, B) U-118 MG at 2% O2, two plates. (C) Astrocytes at 2% O2, one plate. (D) U-118 MG under normoxia, one plate. (E, F) Astrocytes under normoxia, two plates. The color scale is capped at 50 points. The outlined cell marks 4 mM 2-DG plus 2 mM metformin.

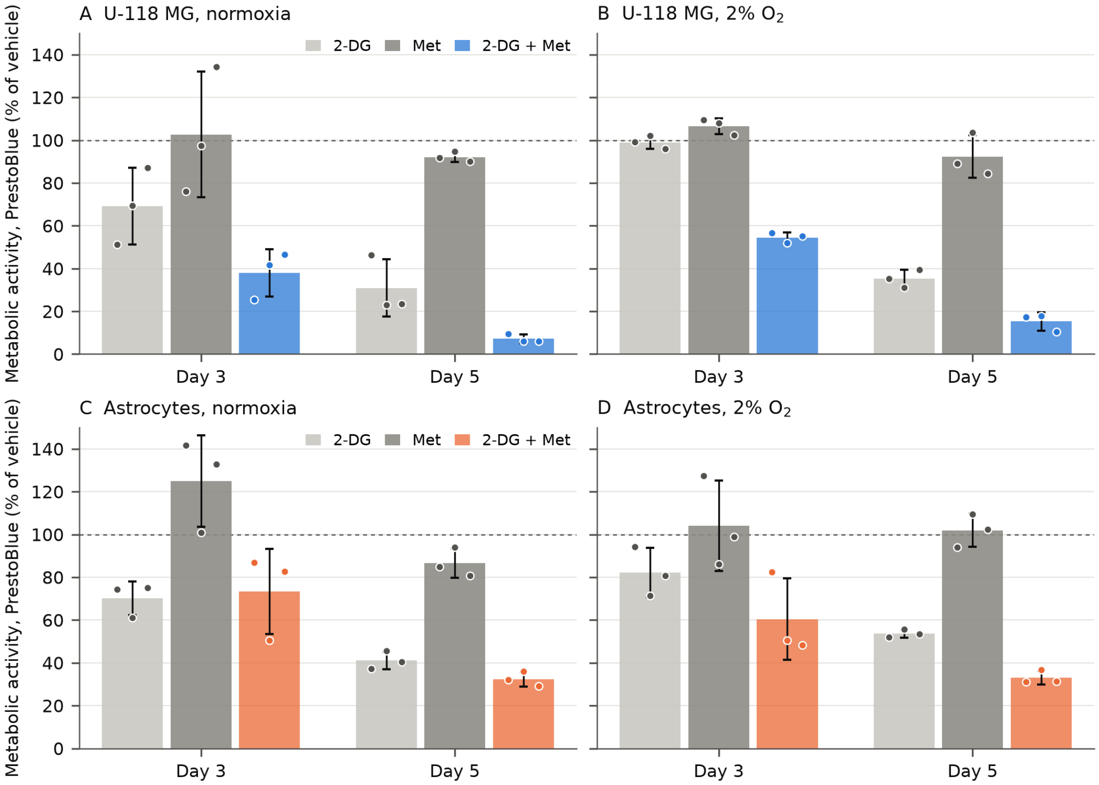


**Figure S3. Metabolic activity by PrestoBlue.** PrestoBlue signal, as percent of the vehicle wells on the same plate, after treatment with 4 mM 2-DG, 2 mM metformin (Met), or both. (A, B) U-118 MG under normoxia and 2% O2. (C, D) Astrocytes under normoxia and 2% O2. The day 3 and day 5 groups are separate experiments, as in Figure 2. Bars show the mean of plates, points the individual plates, error bars SD. n = 3 plates per group. The dashed line marks vehicle (100%).

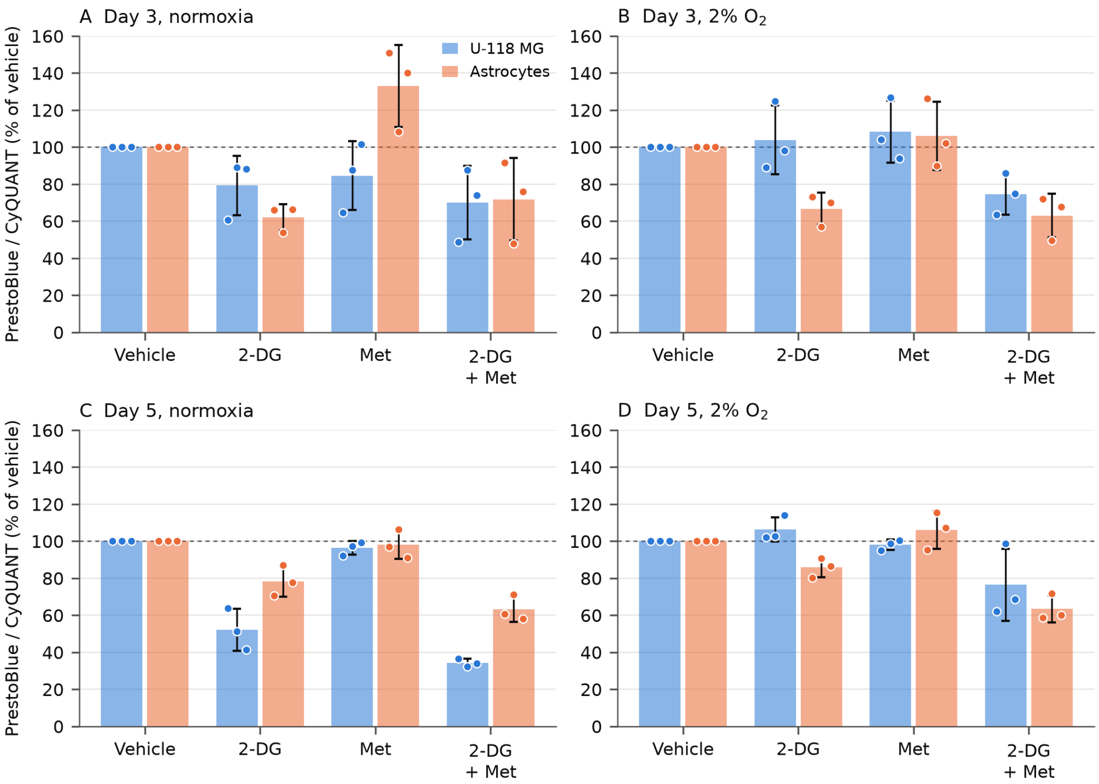


**Figure S4. Metabolic activity per cell.** PrestoBlue divided by CyQUANT, percent of the vehicle wells on the same plate, for U-118 MG and astrocytes treated with vehicle, 4 mM 2-DG, 2 mM metformin (Met), or both. (A, B) Day 3 under normoxia and 2% O2. (C, D) Day 5. Bars show the mean of plates, points the individual plates (independent biological replicates, six wells each), error bars SD. n = 3 plates per group. One vehicle well was excluded at day 5 (U-118 MG, 2% O2, plate 1, well F8). The dashed line marks 100%.
